# Shared etiology of Mendelian and complex disease supports drug discovery

**DOI:** 10.1101/2023.07.23.550190

**Authors:** Panagiotis N. Lalagkas, Rachel D. Melamed

## Abstract

Drugs targeting disease causal genes are more likely to succeed for that disease. However, complex disease causal genes are not always clear. In contrast, Mendelian disease causal genes are well-known and druggable. Here, we seek an approach to exploit the well characterized biology of Mendelian diseases for complex disease drug discovery, by leveraging evidence of pathogenic processes shared between monogenic and complex disease. One way to find shared disease etiology is clinical association: some Mendelian diseases are known to predispose patients to specific complex diseases (comorbidity). Previous studies link this comorbidity to pleiotropic effects of the Mendelian disease causal genes on the complex disease. In previous work studying incidence of 90 Mendelian and 65 complex diseases, we found 2,908 pairs of clinically associated (comorbid) diseases. Using this clinical signal, we can match each complex disease to a set of Mendelian disease causal genes. We hypothesize that the drugs targeting these genes are potential candidate drugs for the complex disease. Our analysis shows that the candidate drugs are enriched among currently investigated or indicated drugs for the relevant complex diseases (odds ratio=1.84, p=5.98e-22). By combining comorbidity with genetic similarity, we recommend drugs further enriched for those investigated or indicated. Our findings suggest a novel way to take advantage of the rich knowledge about Mendelian disease biology to improve treatment of complex diseases.

## Introduction

Traditional drug development pipeline is costly and slow. It is estimated that around $2.6 billion is spent and approximately 12-15 years are required for just one new drug to reach the market^1,2^. Additionally, clinical trial success rates remain low^3^. Therefore, there is a pressing need for new approaches to predict which drugs will succeed.

Recently, genetics has emerged as a resource for predicting drug success. Genome-wide association studies (GWAS) have identified genetic variants associated with complex diseases that are also known therapeutic drug targets^4^. For instance, mutations in the *IL23R* locus have been associated with Crohn’s disease^5^. Ustekinumab, a monoclonal antibody originally approved for the treatment of psoriasis, targets the *IL23* p-40 subunit^6^. Based on genetics, ustekinumab was successfully repurposed for Crohn’s disease^7–9^. This example highlights the importance of genetics both in target prioritization and drug discovery.

Nelson et al. analyzed historical data of clinical trials and found that drug uses supported by human genetic evidence are twice as likely to succeed in clinical trials^10^. Four years later, King et al. confirmed these findings by analyzing data not available at the time of the Nelson et al. study^11^. Moreover, King et al. found that the success rate of a drug is higher when the disease causal gene is clearly identified, as in the case of monogenic (Mendelian) diseases. However, identifying the disease causal genes from GWAS can be challenging as the majority of the GWAS hits are in non-coding regions^12^. In contrast, in monogenic (Mendelian) diseases, the causal genes are both well-known and druggable^13^. Then, developing a way to translate knowledge about Mendelian disease biology to complex diseases could have a significant impact on their treatment.

We have previously exploited clinical data to discover associations between Mendelian and complex diseases^14,15^. In a systematic analysis of 90 Mendelian and 65 complex diseases, Blair et al. used health records to identify which complex diseases individuals with a Mendelian disease are predisposed to, finding 2,908 clinically associated (comorbid) pairs of Mendelian and complex diseases^14^. That study showed evidence that comorbidity can be tied to pleiotropic effects of disease genes. In a follow-up study focusing on cancers, Melamed et al. showed that Mendelian disease causal genes are likely to be frequently mutated in comorbid cancers^15^. For instance, patients with Rubinstein Taybi syndrome, a Mendelian disease caused by mutations in *CREBBP*^16^, are predisposed to lymphoma^17^. *CREBBP* is one of the most frequently inactivated genes in lymphoma^18^ meaning that the observed comorbidity can be attributed to a pleiotropic effect of *CREBBP* mutation causing Rubinstein Taybi syndrome and contributing to lymphoma.

Both of the above studies systematically demonstrate that Mendelian disease comorbidity can suggest a role of Mendelian causal genes on complex disease. However, Mendelian disease comorbidity has not been previously used for complex disease drug discovery purposes. Building on previous findings, we hypothesize that if the Mendelian disease causal genes contribute to the development of a comorbid complex disease, then these genes can be novel therapeutic targets for that disease (***Figure 1A***).

**Figure 1.**
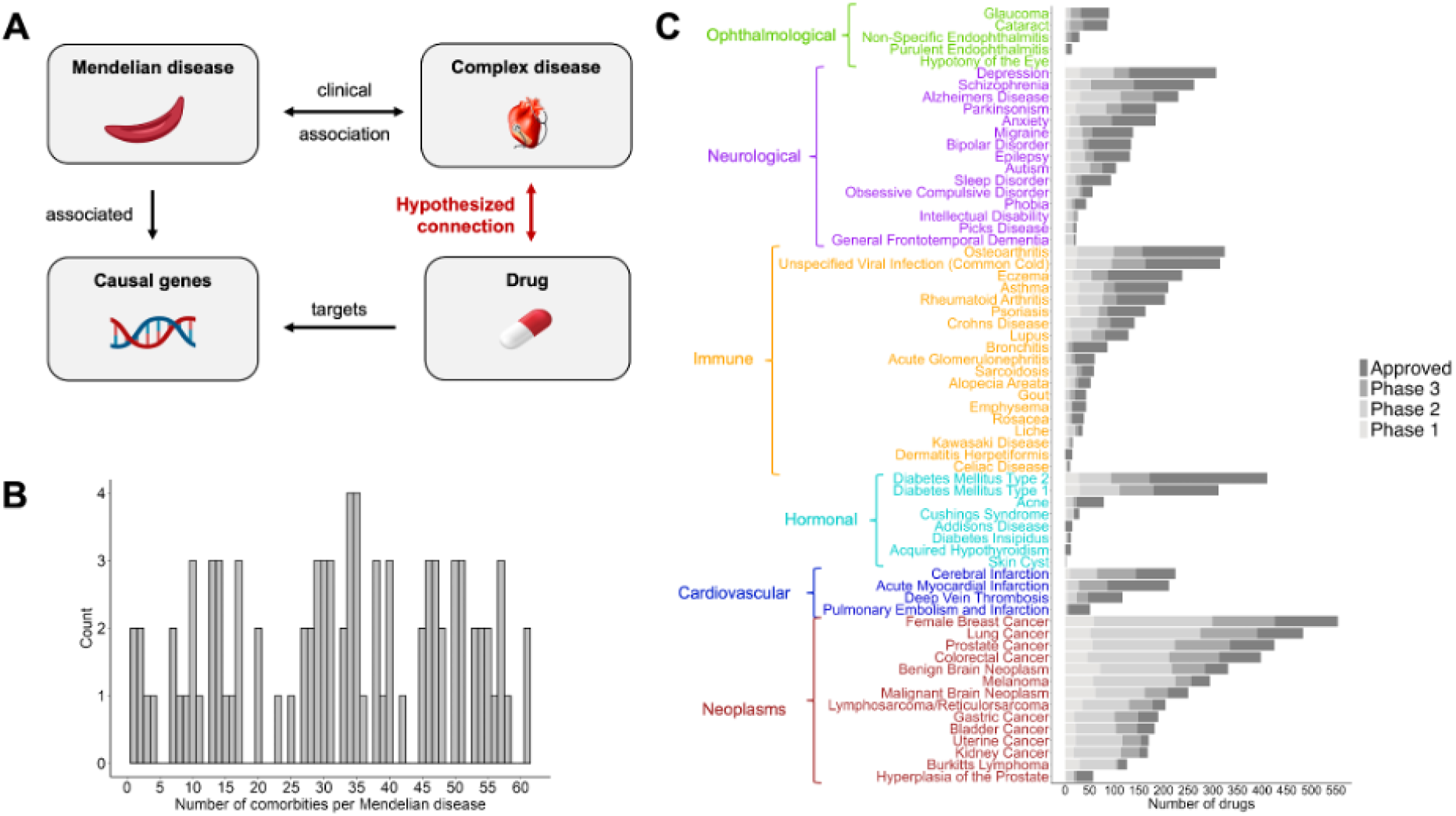
Outline of the approach. **A**. Proposed method where the drugs targeting genes causally associated with a Mendelian disease are suggested as candidate drugs for its clinically associated (comorbid) complex disease. This hypothesized connection between the drug and the complex disease is based on the previously shown pleiotropic effects of the Mendelian disease causal genes on the development of the comorbid complex disease. **B**. Distribution of the number of comorbid complex diseases per Mendelian disease. **C**. Number of investigated (per clinical trial phase) and approved drugs for each complex disease.

## Results

### Integrating data to test Mendelian diseases as a resource for drug repurposing

From Blair et al.^14^, we obtain clinical associations between 2,908 pairs of a Mendelian and a complex disease. The data include 90 Mendelian diseases with known causal genes and 65 complex diseases across six disease categories (cardiovascular, hormonal, immune, neoplasms, neurological, ophthalmological). ***Figure 1B*** shows the distribution of the number of comorbid complex diseases per Mendelian disease. Using drug-gene target information from DrugBank and Mendelian causal genes from the Online Mendelian Inheritance in Man (OMIM), we compile a list of 781 drugs that target the Mendelian disease causal genes. This allows us to suggest candidate drugs for each complex disease based on its comorbid Mendelian diseases (***Figure 1A***).

To test our hypothesis, we compare our candidate drugs against drugs currently investigated or indicated for the complex diseases. We curate 29,758 clinical trials that investigate 1,795 drugs for 64 complex diseases (median of 110 investigated drugs per complex disease). In addition to the investigated drugs, we compile current approved drug uses, including 1,373 indicated drugs for 64 complex diseases (median of 42 indicated drugs per complex disease) (***Figure 1C***).

### Mendelian disease comorbidity identifies drugs under current investigation or indication

First, we assess whether the candidate drugs for a complex disease are enriched for those currently investigated or indicated for that disease. Accounting for the number of gene-targets per drug and the disease category, we find that the candidate drugs are significantly enriched for drugs currently investigated or indicated (odds ratio=1.84, p=5.98e-22) (***Figure 2A***).

**Figure 2.**
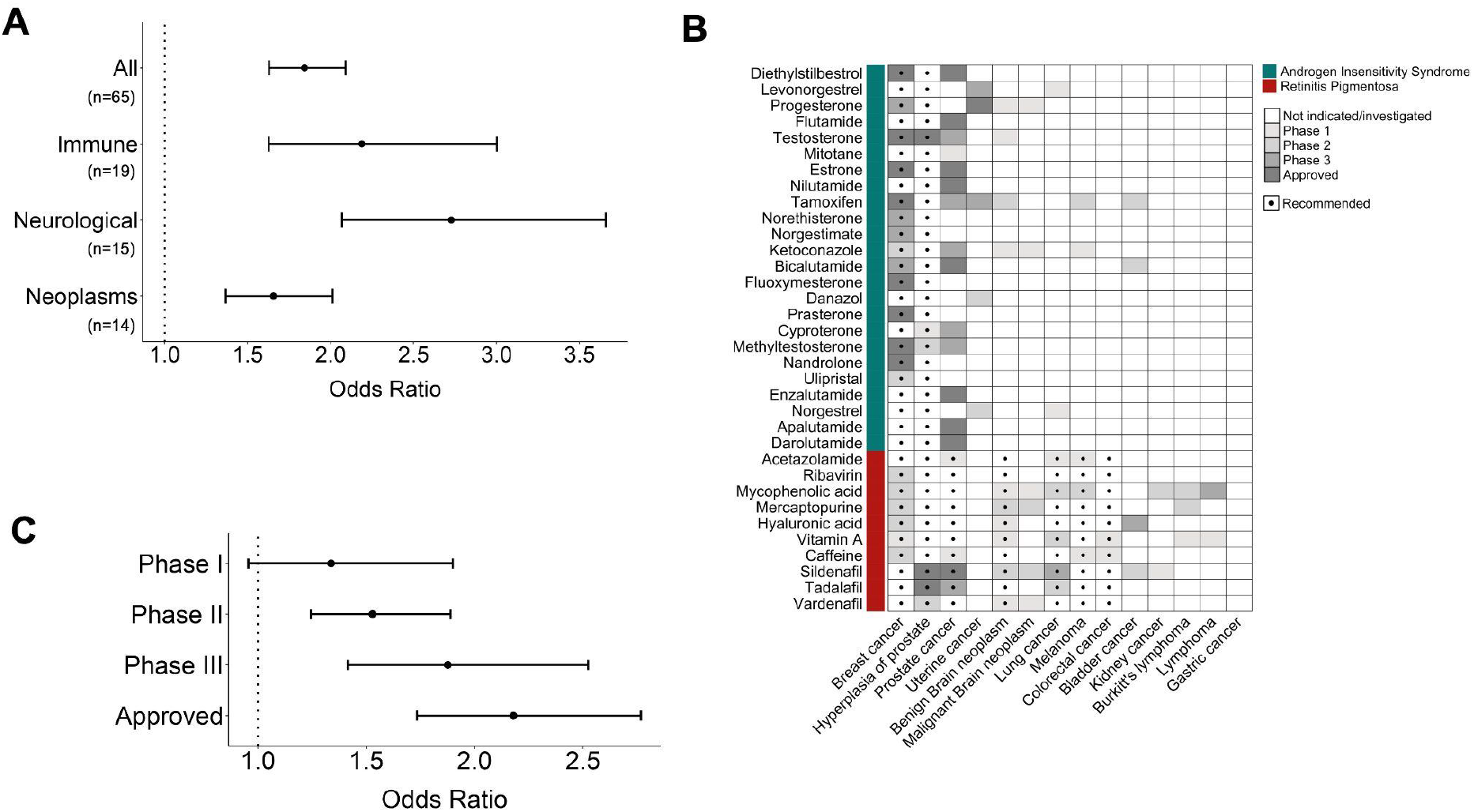
Clinical associations between Mendelian and complex diseases predict candidate drugs with higher potential of success for the complex diseases. **A**. Odds ratio of candidate drugs to be currently investigated or indicated for the complex diseases within a disease category. Only disease categories that are significant compared to 1,000 permutations of the comorbidity relationships (p_permutation_<0.05) are shown. **B**. Examples of recommended candidate drugs for 14 neoplasms based on their clinical associations with 2 Mendelian diseases: Androgen Insensitivity Syndrome and Retinitis Pigmentosa. Gray-scaled boxes indicate the phase of the drug in the development pipeline for each neoplasm. **C**. Odds ratio of candidate drugs to be investigated or indicated for a complex disease per drug development phase. *in A and C, bars represent the observed odds ratio with 95% confidence intervals.

Next, we seek to exclude artifactual explanations for this signal. One such artifact is variation in disease frequency: disease frequency can impact both the ability to discover disease genes and the power to discover clinical associations. To exclude this spurious source of association, we randomly permute which complex diseases each Mendelian disease is comorbid with. This random permutation preserves the characteristics of each Mendelian disease, such as number of complex disease comorbidities, but not the list of candidate drugs for each complex disease. After 1,000 permutations, we find that the observed association is significantly stronger than expected by chance (p_permutation_<0.001). For Mendelian diseases with many comorbidities, the permutation does not impact recommendations; therefore, the permuted signal, though weaker, still has odds ratio > 1. However, when these Mendelian diseases are removed, comorbidity is significantly predictive of drug uses, while the permuted comorbidity is not predictive (***Figure S1***).

We repeat the analysis at a disease category level, and we find significant results for neurological, immune, neoplasms, ophthalmological, and hormonal diseases. However, after permutation analyses, only neurological, immune, and neoplasm disease categories remain significant (p_permutation_<0.05) (***Figures 2A, S2-S7***). This may be due to the low number of analyzed complex diseases that fall under the cardiovascular (n=4), ophthalmological (n=5), and hormonal (n=9) disease categories, potentially reducing the statistical power to detect a significant association. Additionally, these disease categories have a lower number of current therapies (***Figure 1C***). ***Figure 2B*** shows an example of our recommended candidate drugs for 14 neoplasms, illustrating an extensive overlap between the candidate drugs and the drugs currently investigated or indicated for these neoplasms.

Next, we ask if the candidate drugs are more likely to be in advanced drug development phases for the relevant complex diseases. To test this, we stratify drugs by their drug development phase for a complex disease: phase I, phase II, phase III, indicated. First, we find a significant enrichment of candidate drugs for a complex disease among drugs in any of phase I, II, or III (odds ratio=1.59, p=2.27e-09; p_permutation_<0.001) (***Figure S8***). Stratified per clinical trial phase (phase I, II, or III), we find a progressive increase in the enrichment for drug success with increasing phase (p_permutation_<0.05) (***Figures 2C, S9-11***). Additionally, when considering only indicated drugs for a complex disease, we find an even greater enrichment of its candidate drugs for drug success (odds ratio=2.18, p=5.94e-11; p_permutation_<0.001) (***Figures 2C, S12***). Overall, our predicted drug candidates show more enrichment in categories with more clinical evidence, supporting the potential of our approach for identifying new successful drugs.

### Prioritizing Mendelian diseases targeted by high number of drugs

Mendelian disease causal genes are known to be good drug targets^14^. We find 193 out of 593 Mendelian genes (32.6%) to be targeted by at least one drug (median: 2 drugs per Mendelian gene). However, outliers exist: androgen receptor (*AR*), a gene mutated in Androgen Insensitivity Syndrome, is targeted by 82 drugs^19^. This variation in drug targeting of Mendelian genes may suggest that certain disease processes are more druggable. We hypothesize that the most druggable Mendelian diseases are the most promising for providing insight into complex disease therapeutics.

To test this hypothesis, we repeat the above analysis for each Mendelian disease individually, for Mendelian diseases targeted by at least one drug (n=68) (***Figure 3A***). That is, we test whether the drugs targeting the causal genes of each Mendelian disease are enriched for drugs currently investigated or indicated for its comorbid complex diseases. Although testing only the drugs targeting a single Mendelian disease reduces the statistical power of the analysis, we find 8 significant Mendelian diseases (p_permutation_<0.05). Further, we find that these 8 Mendelian diseases are targeted by a significantly higher number of drugs than other Mendelian diseases (p=9.1e-05, one-sided Wilcoxon rank-sum test) (***Figure 3B***). To exclude the possibility that this is due only to higher numbers of drugs increasing power to discover an association, we compare the result against a permutation analysis that permutes the drugs targeting each Mendelian disease (p_permutation_=0.018).

**Figure 3.**
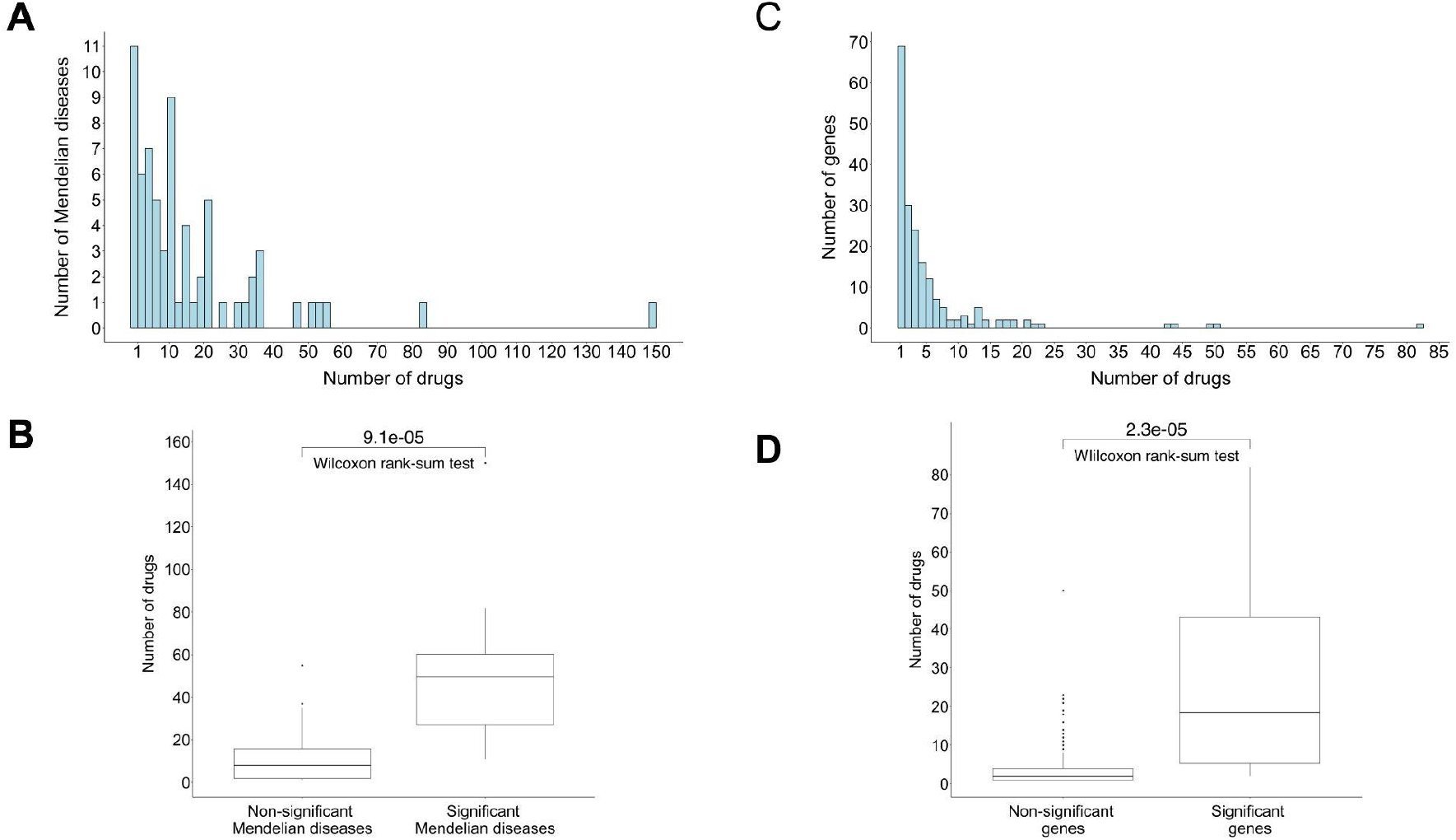
Highly drugged Mendelian diseases are a better resource for candidate drugs. **A**. Histogram of the number of drugs targeting a Mendelian disease, for Mendelian diseases targeted by at least one drug (n=68). **B**. Mendelian diseases that significantly predict candidate drugs already investigated or indicated for their comorbid complex diseases are targeted by a higher number of drugs compared to the other Mendelian diseases (p=9.1e-05, one-sided Wilcoxon rank-sum test comparing the number of drugs in each group). **C**. Histogram of number of drugs targeting a Mendelian gene, for genes targeted by at least one drug (n=193). **D**. Genes linked to Mendelian diseases that significantly predict candidate drugs already investigated or indicated for their comorbid complex diseases are targeted by a higher number of drugs compared to the other genes (p=2.3e-05, one-sided Wilcoxon rank-sum test comparing the number of drugs in each group).

In another test of this hypothesis, we ask which Mendelian disease genes successfully point to new drug indications. That is, for each Mendelian disease gene, we use comorbidity to suggest which complex diseases may benefit from drugs targeting that gene. Under our hypothesis, we expect that highly druggable genes can more successfully be used for finding new drug uses. To test this, we repeat the above analysis for each gene targeted by at least one drug (n=193) (***Figure 3C***), comparing the association to permutations. We find 12 significant genes (p_permutation_<0.05), and these successful genes are again targeted by a higher number of drugs compared to the other Mendelian disease genes (p=2.3e-05, one-sided Wilcoxon rank-sum test) (***Figure 3D***). Altogether, these results imply that Mendelian diseases associated with more druggable genes are a particularly promising resource for complex disease therapeutics.

### Combining comorbidity with genetic similarity enhances drug predictions

Comorbidity is a way to discover diseases sharing a biological basis, but it is not the only way. Comorbid Mendelian and complex diseases have been shown to be more likely to share related or overlapping genes, which is known as genetic similarity^14,15^. Additionally, genetic similarity between drug targets and disease-linked genes has also been shown to predict successful drugs for a disease^10,11^. Building on these results, we propose that genetic similarity could contribute to discovering therapeutically relevant shared etiology of Mendelian and complex diseases^20,21^. Specifically, we propose that by combining comorbidity with genetic similarity, the two forms of evidence can more robustly point to diseases with shared etiology, increasing the predictive success of our approach.

To test this hypothesis, we focus on cancers, one of the disease categories with the strongest association in our analysis (***Figure 2A***). Cancers are also of interest because each type of cancer has been associated with a set of recurrently mutated driver genes in The Cancer Genome Atlas (TCGA); we previously showed that Mendelian diseases comorbid with a cancer are enriched for genetic similarity to somatically mutated cancer driver genes^15^. Building on that work, we ask whether candidate drugs supported by both comorbidity and genetic similarity between Mendelian disease and cancer have greater probability for success. Among the 10 cancers in TCGA, Mendelian disease comorbidity again predicts drugs enriched for those currently investigated or indicated (odds ratio=1.69, p=7.42e-06, p_permutation_=0.014) (***Figure S13***). But, combining comorbidity with genetic similarity, drugs with both forms of evidence are even more enriched for drugs with clinical support (odds ratio=2.19, p=6.33e-13, p_permutation_=0.001) (***Figures 4A, S14***).

**Figure 4.**
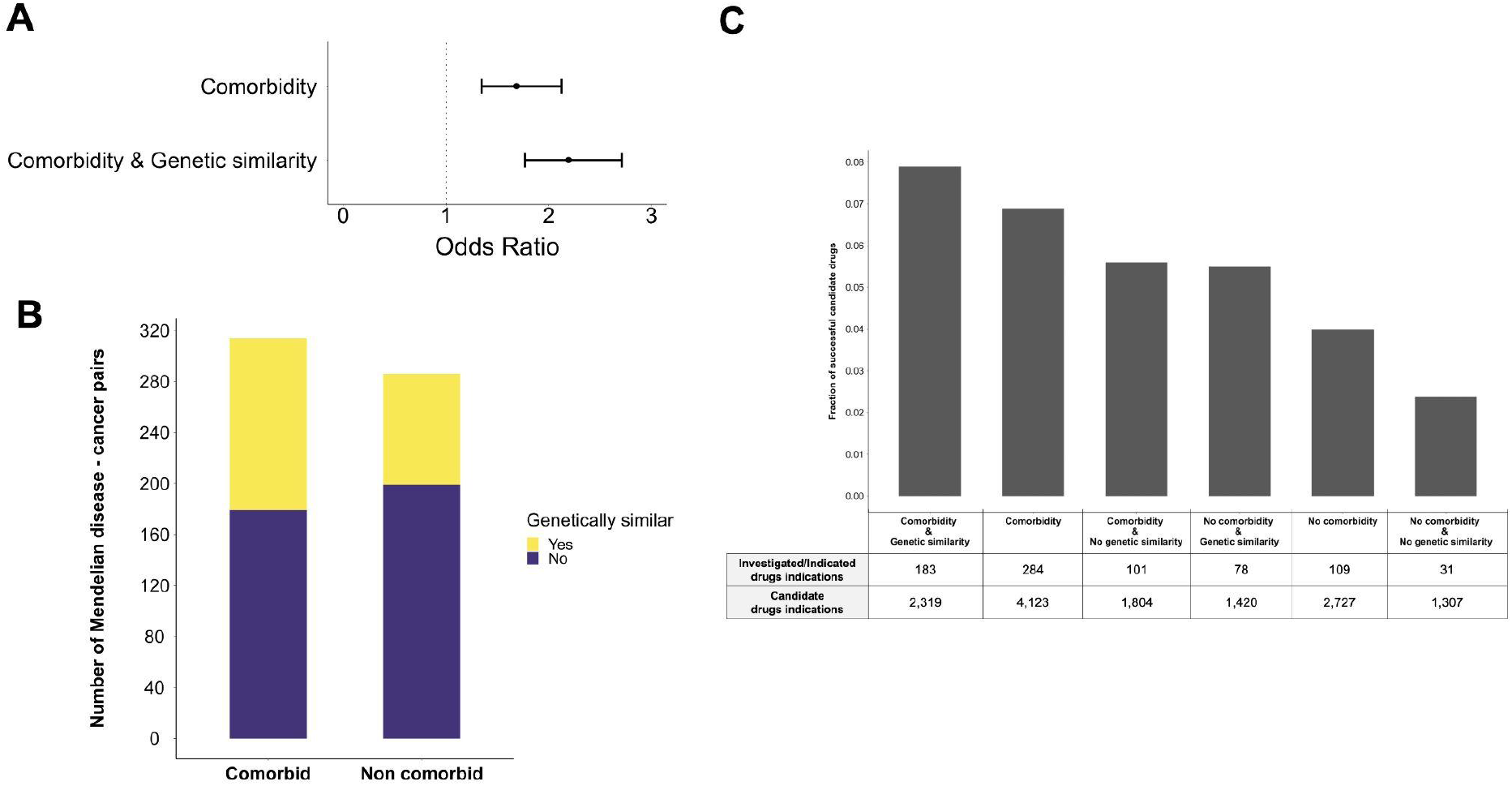
Combination of comorbidity and genetic similarity prioritizes candidate drugs for cancers with higher probability of success. **A**. Odds ratio of success for candidate drugs supported by comorbidity or comorbidity and genetic similarity between Mendelian disease and cancer. Information about genetic similarity comes from Melamed et al.^15^. Black bars represent the 95% confidence intervals. **B**. Number of comorbid and genetically similar pairs among 60 Mendelian diseases and 10 cancers. Genetic similarity was estimated using two metrics established here: gene overlap and co-expression. **C**. Percentage of candidate drug-cancer pairs to be currently investigated or indicated among different levels of support.

To investigate the contributions of genetic similarity and comorbidity individually and combined, we stratify the 600 pairs of 60 Mendelian diseases and 10 cancers into those that are comorbid and those with no detected comorbidity relationship. As genetic similarity was not previously evaluated for non-comorbid disease pairs, we establish two measures for genetic similarity between two diseases, gene overlap and gene co-expression, similar to the measures used in Melamed, et al.^15^ (see Methods). Among 314 comorbid disease pairs, 135 are also genetically similar (43%). These comorbid and genetically similar pairs greatly overlap with the ones identified by Melamed et al.^15^ (p=7.13e-08, one-sided Fisher’s exact test), indicating that our genetic similarity metrics are consistent with the prior work. Among the remaining 286 non-comorbid pairs, 87 are genetically similar (30.4%) (***Figure 4B***). The higher rate of genetic similarity among the comorbid diseases is consistent with the prior literature^15^.

Using all the drugs targeting the causal genes of the 60 Mendelian diseases, we compile a list of 6,850 possible drug-cancer pairs (685 drugs x 10 cancers). Among 2,727 drug-cancer pairs not supported by comorbidity, we find that those supported by genetic similarity have increased probability of drug success (odds ratio=2.32, p=1.07e-04). This implies that genetic similarity might be able to detect shared etiology between Mendelian disease and cancer pairs that cannot be detected with comorbidity. Further, among 4,123 drug-cancer pairs supported by comorbidity, those additionally supported by genetic similarity have greater probability of drug success (odds ratio=1.39, p=0.01). As we expect that candidate drug recommendations supported by comorbidity are already enriched for shared etiology, it is logical that the effect of genetic similarity would be smaller for this category of recommendations, but the effect is still significant. Notably, drug uses supported by both comorbidity and genetic similarity are most enriched for known drug uses (***Figure 4C***, most left bar).

In conclusion, these findings suggest that by combining the two forms of evidence we can prioritize candidate drugs that target the shared biology between two comorbid diseases, enhancing the use of Mendelian disease biology for drug discovery.

## Discussion

Previous studies have suggested that Mendelian disease genes pleiotropically contribute to the development of complex diseases, resulting in significantly increased risk of the complex disease in individuals with the Mendelian disease^14,15^. However, this insight has not been harnessed for drug discovery. Here, we have shown that comorbidity between Mendelian and complex diseases can recommend candidate drugs for the complex diseases. Importantly, these candidate drugs are more likely to be in advanced drug development phases or have received regulatory approval, suggesting that Mendelian disease comorbidity can be used to prioritize drugs with high potential of eventual approval.

Our findings provide a novel way to leverage the well-known biology of Mendelian diseases to enhance the treatment of complex diseases. For instance, verapamil, an approved calcium channel inhibitor for the treatment of angina^22^, is among our recommended candidate drugs for Type 1 Diabetes (T1D). This recommendation is supported by the comorbidities of T1D with Long QT Syndrome (*CACNA1C*) and Spinocerebellar Ataxia (*CACNA1A*). Studies in mice have previously demonstrated verapamil’s potential to prompt the survival of insulin-producing β-cells and reverse T1D^23^. Notably, verapamil has recently been tested in a phase III clinical trial for T1D treatment^24^. Additionally, we recommend carbamazepine, an approved sodium channel inhibitor for the control of seizures^25^, as a candidate drug for the treatment of T1D based on its comorbidities with Long QT Syndrome (*SCN5A*) and Erythromelalgia (*SCN9A*). This recommendation is further supported by preclinical studies showing that inhibition of sodium channels increases the expression of *INS1* and *INS2* and thus protects from the development of T1D^26–28^. Looking ahead, we anticipate that future clinical trials should consider testing the efficacy of this drug category for preventing T1D.

We also present an approach for identifying a subset of Mendelian diseases with the most utility for drug discovery. In general, Mendelian diseases are enriched for drugged genes^13^, but some Mendelian diseases appear to be targeted by even more drugs than the average. Focusing on both the Mendelian disease and gene level, we find that diseases associated with highly drugged genes hold greater promise for future drug discovery efforts.

As well, building on previous work that prioritizes drugs functionally related to disease genes^15^, we explore genetic similarity as an additional way to identify diseases sharing likely pleiotropic causal genes. In an analysis of ten cancers, we find that candidate drugs supported by both comorbidity and genetic similarity between a Mendelian disease and a cancer have greater probability of success. By combining two independent sources of evidence for shared disease etiology, future research can use Mendelian disease genes to prioritize new drug uses.

Our work has some limitations. First, we could not compile a complete list of investigated drugs for the 65 complex diseases due to annotation inconsistencies. Second, the complete list of genes causally associated with a Mendelian disease might not be complete due to its rarity. Third, comorbidity may not always be due to pleiotropic effects of the Mendelian disease genes on the development of the complex diseases, but it can also be due to indirect or interaction effects. Similarly, lack of measurable comorbidity between a pair of diseases does not definitively mean an absence of shared pathological processes but could be due to disease frequency or interaction effects. Finally, we find that the enrichment of candidate drugs for success varies across disease categories: our results were not significantly predictive for cardiovascular, ophthalmological, and hormonal diseases. This may be because we were not able to test a diverse set of diseases in these categories leading to reduced statistical power.

In conclusion, we leverage the well-known biology of Mendelian diseases to improve treatment of common diseases. To our knowledge, this is the first study that suggests the use of clinical associations of Mendelian diseases to inform drug discovery. Future work can both exploit the drugs we suggest for each disease and explore Mendelian disease genes currently lacking drugs as novel drug targets. In fact, according to Finan et al.^29^, almost one fourth (24.4%) of the undrugged Mendelian genes, have high druggability potential. Additionally, disease comorbidity might improve other drug repurposing efforts when considered as an additional source of evidence for prioritizing drug repurposing candidates. Finally, future efforts can also build on the idea by investigating whether clinical associations between common diseases can expand the use of existing drugs.

## Methods

### Recommending drugs for complex disease based on Mendelian disease comorbidities

We download comorbidity pairs between 95 Mendelian and 65 complex diseases from the Blair et al. supplementary materials^14^. We remove 5 Mendelian diseases (and their comorbidities) that are due to chromosomal abnormalities and the causal gene is not obvious (Down Syndrome, Edward Syndrome, Klinefelter Syndrome, Patau Syndrome, Turner Syndrome). We use the remaining 2,908 comorbidity pairs between 90 Mendelian diseases and 65 complex diseases in our main analysis. Additionally, we group the 65 complex diseases into 6 major disease categories: cardiovascular (n=4), hormonal (n=8), immune (n=19), neoplasms (n=14), neurological (n=15), ophthalmological (n=5).

We systematically code all the 65 complex diseases using MeSH codes. To do this, we download the supplementary “Table S2” from Blair et al.^14^ which contains the ICD-10 billings codes the authors used to identify the complex diseases. After manual matching to relevant MeSH codes, we have 230 unique MeSH codes for 65 complex diseases (median of 3 MeSH codes per complex disease).

We download information for 5,800 drugs from DrugBank (version 5.19, date of download: June 16, 2022). After filtering for gene targets in humans, for each drug, we keep its DrugBank ID and its gene targets (HGNC symbols). Drugs included in this file can be approved, investigational, small molecule, biotech, experimental, nutraceutical, illicit, or withdrawn. We do not filter the list of drug-gene targets based on pharmacological action.

To suggest drug repurposing candidates for a complex disease, we first find its comorbid Mendelian diseases. We obtain the genes causally associated with these Mendelian diseases from the OMIM. Using drug-gene target information from DrugBank, we find drugs targeting the Mendelian disease causal genes and we suggest them as candidate drugs for the complex disease.

### Finding investigated drugs for the complex diseases

To find drugs currently investigated for the 65 complex diseases in our sample, we download data for 432,597 clinical trials from the Aggregate Content of ClinicalTrials.gov (AACT) database in a pipe-delimited format (data of download: November 4, 2022)^30^. For each clinical trial, we keep the clinical trial ID, clinical trial phase, conditions (diseases) studied and interventions (drugs) tested. We filter out clinical trials tested behavioral, device, diagnostic test, dietary supplements, procedures, radiation, or “other interventions”, and do not provide MeSH terms for both conditions and interventions. For the total of 109,430 remaining clinical trials, we match the MeSH terms of both conditions and interventions to MeSH codes using the Unified Medical Language System (UMLS) database. We group clinical trial phases to Phase I (Phase I and Early Phase I), Phase II (Phase II and Phase I/Phase II), Phase III (Phase III and Phase II/Phase III) or unknown phase (no information provided). Phase IV studies are conducted after a drug gets approved to find long-term benefits and side-effects that could not be discovered in the duration of a clinical trial. Therefore, we consider drugs in Phase IV clinical trials as indicated drugs (see below). Eventually, we have 31,053 clinical trials that tested interventions for the 65 complex diseases in our sample.

For these clinical trials, we convert the MeSH codes of interventions to DrugBank IDs using the UMLS API (crosswalk function). However, DrugBank does not assign IDs to drug combinations. To include them in our analysis, we convert the MeSH codes that did not match to a DrugBank ID, to RxNORM CUIs using the UMLS API (crosswalk function). Then, for each drug combination, we obtain each active pharmaceutical ingredient using the UMLS API (“Retrieving Source-Asserted Relations”; vocabulary=RXNORM; relation label=has_part). Moreover, MeSH vocabulary assigns different codes to each form of an active pharmaceutical ingredient. But DrugBank assigns IDs only to the general forms. For example, liposomal doxorubicin and doxorubicin are two separate entries in MeSH vocabulary but not in DrugBank (doxorubicin). To deal with this discrepancy, we follow the steps described above and we obtain the active pharmaceutical ingredients using the UMLS API (“Retrieving Source-Asserted Relations”; vocabulary=RXNORM; relation label=form_of). Then, for each drug, we convert its RxNORM CUI to DrugBank ID using the UMLS API (crosswalk function).

Eventually, we have 29,758 clinical trials that tested 1,795 drugs for 64 complex diseases in our sample. Note that the complex disease “Dermatitis herpetiformis” did not have any investigated drugs at the time of this study.

### Finding indicated drugs for the complex diseases

To find drugs that are currently indicated for the 65 complex diseases, we download 4,225 approved drugs from DrugBank (version 5.19, date of download: June 16, 2022). We get their indications by combining information from RxNORM and repoDB, as described below.

Using the RxNORM API (getClassByRxNormDrugName function), we obtain diseases (in MeSH terms) with a relationship of “may_treat” or “may_prevent” with each approved drug.

We then match the diseases to MeSH codes using the UMLS database.

repoDB is a publicly available database that contains drug repositioning successes and failures by integrating data from DrugCentral and ClinicalTrials.gov^31^. We download the full database and, for each drug, we keep its DrugBank ID and approved indication(s), after excluding the ones with a note of suspended, terminated, or withdrawn. All indications are coded in UMLS CUIs, so we convert them to MeSH codes using the UMLS database.

After combining the data from RxNORM and repoDB, we have 939 unique drugs indicated for 58 complex diseases. We then add to this data set the drugs in clinical trials Phase IV to get a total of 1,373 unique drugs indicated for 64 complex diseases. Note that the complex disease “hypotony of the eye” did not have any indicated drug at the time of this study.

### Statistical analysis

#### Logistic regression to evaluate candidate drugs

We find 781 unique drugs that target the causal genes of the 90 Mendelian diseases. Using these drugs and the 65 complex diseases, we create a table where each row is a drug-complex disease pair. Therefore, in our main analysis, the number of rows in this table is 50,765 (781 drugs multiplied by 65 complex diseases). We assess whether our recommended drug-disease pairs are predictive of current investigated or indicated drug-disease pairs in a logistic regression model also adjusting for i) the disease category of each complex disease; ii) the number of known gene-targets per drug. We account for the disease category due to differences in the number of drugs investigated or indicated among the 6 disease categories tested in this study (***Figure 1B***). Additionally, we account for the number of targets per drug as drugs with a higher number of known targets are more likely to be linked to a Mendelian disease and may be more likely to be subject to research investment for new indications.

To ensure that class imbalance does not bias the coefficient estimate in our model, we also conducted a weighted logistic regression. As shown in **Figure S15**, the results from the weighted logistic regression are also significant and comparable to the non-weighted logistic regression. Therefore, we use a non-weighted logistic regression in all of our analyses.

#### Permutation tests to assess the significance of the observed associations

In our main analysis, we want to assess the significance of the observed associations between drug repurposing candidates and currently investigated or indicated drugs for a complex disease. The random permutation changes which complex diseases are comorbid with each Mendelian disease, by keeping unchanged all the intrinsic Mendelian disease characteristics, such as prevalence, number of comorbidities, and associated causal genes. This allows us to assess if the observed association is solely attributed to the Mendelian disease comorbidity or not.

In the per Mendelian disease analysis, we want to assess if the higher druggability of a Mendelian disease gene rather than the information about comorbidity drives the results. The random permutation changes which drugs target a Mendelian disease gene by keeping unchanged the total number of drugs targeting the Mendelian disease gene.

In both cases, we perform 1,000 permutations to create a null distribution of odds ratios using the logistic regression model above. We then compare the observed odds ratio to this null distribution. We calculate the probability of observing an odds ratio at least as extreme as the original one by estimating the number of times a permuted odds ratio is higher or equal to the observed odds ratio (odds_ratio_observed_ ≤ odds_ratio_permutation_). A result is considered significant if the calculated probability is less than 0.05 (p_permutation_<50/1000).

### Genetic similarity between Mendelian diseases and cancers

We assess the genetic similarity between a Mendelian disease and a cancer using two sources of evidence. First, we consider the extent of genetic overlap between two diseases. This simple metric captures the shared driver genes between two diseases. To capture a wider range of functional relationships between two diseases, we also use gene co-expression across diverse human tissues. A Mendelian disease and cancer pair are considered genetically similar if at least one metric is significant (p<0.05). Both metrics are described below.

The genetic overlap metric tests the significance of the overlap between the Mendelian disease causal genes and the genes significantly altered in a cancer. For each Mendelian disease, we compile a list of causally associated genes using the OMIM database. For each cancer, we compile a list of driver genes using the Broad GDAC Firehose database including genes significantly mutated (as identified by MutSig v2.0, q<0.05) and genes with significant copy number alterations (as identified by Gistic2; q<0.05; peaks with at maximum 50 genes). Then, for a Mendelian disease and cancer pair, we test the significance of the overlap between the set of Mendelian disease causal genes and the set of genes significantly altered in cancer (Fisher’s exact test, p<0.05).

The co-expression metric tests correlation in expression between a cancer driver gene and a set of Mendelian disease genes. To assess genetic similarity using this metric, we first download summarized expression data for 20,162 genes across 37 GTEx tissues from the Human Protein Atlas. We remove 889 genes that do not have expression data across all 37 tissues. The remaining data contained expression for 574 out of the 594 Mendelian disease causal genes. Consequently, the co-expression metric of one Mendelian disease (“Familial Dysautonomia”) with any cancer could not be measured (it was tested using only the genetic overlap metric). The metric tests whether the set of Mendelian disease genes exhibit stronger correlation in expression with the cancer gene compared to the correlation distribution of all other genes with the same cancer gene. We test this using the Wilcoxon rank-sum test and we adjust the resulting p-values to account for the number of cancer genes tested (Benjamini-Hochberg method; p<0.05).

## Code and data availability

Code and publicly available data for reproducing the results and figures presented here are stored on GitHub: https://github.com/lalagkaspn/mendelian_comorbidity_therapeutics.git. Certain datasets and tools require user license. In such cases, links to download the necessary files from the original sources, after obtaining the relevant license, are provided within the scripts.

---

**Figure S15.**
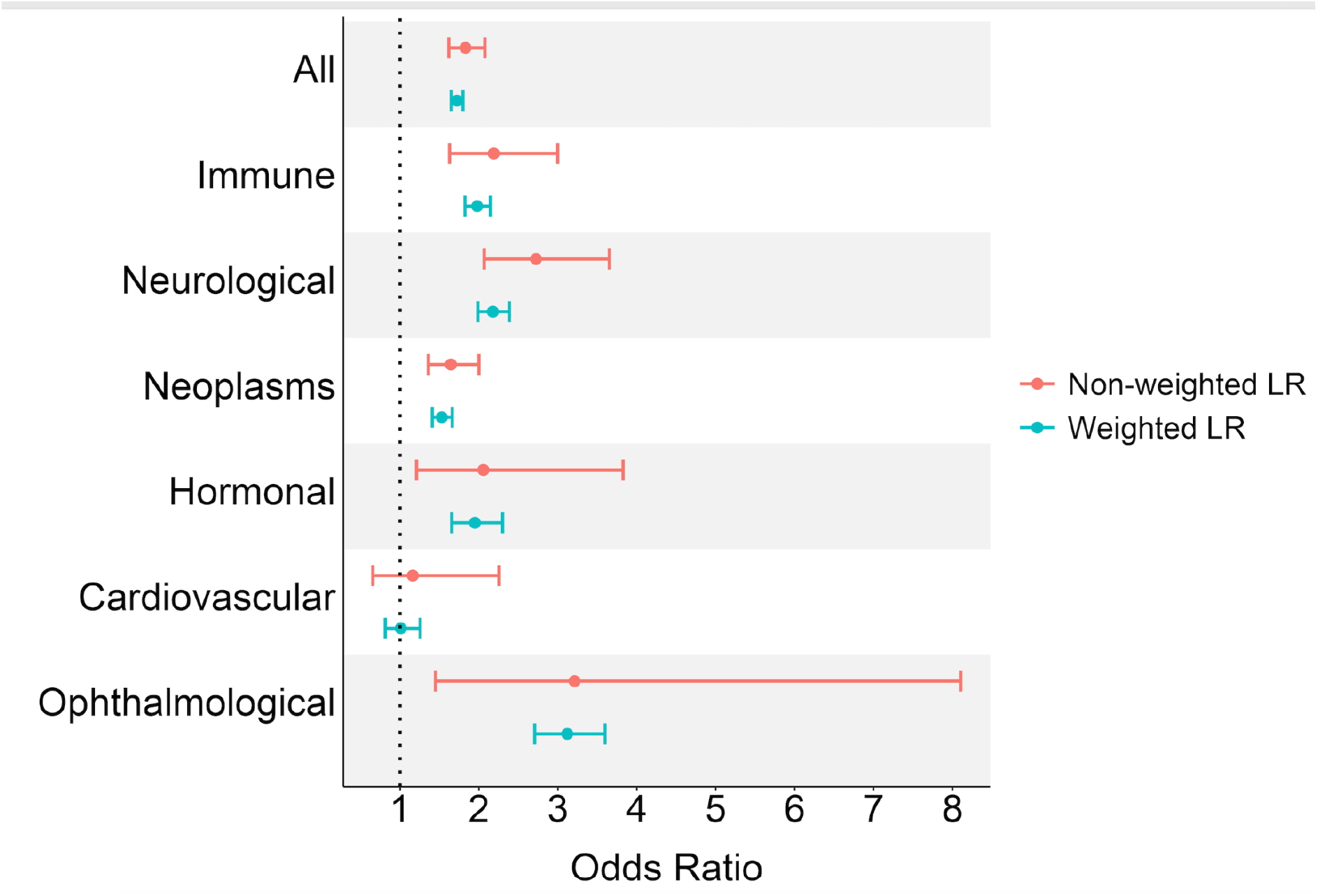
Comparison of odds ratio of recommended drug-disease pairs to be currently investigated or indicated using a non-weighted or weighted logistic regression (LR). Class weights for the weighted LR are estimated using inverse class frequencies, as suggested by King et al (https://gking.harvard.edu/files/0s.pdf). The estimated odds ratio from the weighted LR are again significant and comparable to the estimated odds ratio from the non-weighted LR.

